# Zfp106 binds to G-quadruplex RNAs and inhibits RAN translation and formation of RNA foci caused by G4C2 repeats

**DOI:** 10.1101/2023.03.05.531055

**Authors:** Barbara Celona, Haifan Wu, Bobo Dang, Huong T. Kratochvil, William F. DeGrado, Brian L. Black

## Abstract

Expansion of intronic GGGGCC repeats in the *C9orf72* gene causes amyotrophic lateral sclerosis (ALS) and frontotemporal dementia (FTD). Transcription of the expanded repeats results in the formation of RNA-containing nuclear foci and altered RNA metabolism. In addition, repeat-associated non-AUG (RAN) translation of the expanded GGGGCC-repeat sequence results in the production of highly toxic dipeptide-repeat (DPR) proteins. GGGGCC-repeat-containing transcripts form G-quadruplexes, which are associated with formation of RNA foci and RAN translation. Zfp106, an RNA-binding protein essential for motor neuron survival in mice, suppresses neurotoxicity in a *Drosophila* model of *C9orf72* ALS via a previously unknown mechanism. Here, we show that Zfp106 inhibits formation of RNA foci and significantly reduces RAN translation caused by GGGGCC-repeats in mammalian cells. Further, we show that Zfp106 binds to RNA G-quadruplexes and causes a conformational change in the G-quadruplex structure formed by GGGGCC-repeats. These data suggest that Zfp106 suppresses GGGGCC repeat-mediated cytotoxicity through alteration of the repeat’s G-quadruplex structure.

## INTRODUCTION

Zinc finger protein 106 (Zfp106) is a Cys2-His2 (C2H2) zinc finger protein with four predicted zinc fingers and seven WD40 domains (1-3). Zfp106 primarily localizes to the nucleolus and to nuclear speckles adjacent to spliceosomes (1-3), and previous studies have demonstrated that it functions as an RNA-binding protein with suggested roles in multiple aspects of RNA metabolism, including pre-rRNA processing, splicing, and polyA mRNA binding (1, 2, 4-6). Knockout of the *Zfp106* gene in mice results in profound neuromuscular disease, resembling amyotrophic lateral sclerosis (ALS), with onset at approximately 4-6 weeks of age followed by rapidly progressing motor neuron degeneration and death by 16 weeks (1, 2, 7). Restoration of Zfp106 expression specifically to motor neurons significantly suppresses the phenotype in knockout mice, suggesting that the requirement of Zfp106 for neuromuscular function and motor neuron viability is autonomous to motor neurons (2).

The most common cause of familial ALS and frontotemporal dementia (FTD), a related neurodegenerative disorder of the brain, results from abnormal expansion of the hexanucleotide sequence GGGGCC in the first intron of the *C9orf72* gene (8, 9). Healthy individuals usually have fewer than 30 copies of the hexanucleotide repeat sequence, whereas affected patients may have hundreds to thousands of repeats in their *C9orf72* gene (10). The expanded GGGGCC repeats have been proposed to cause neurodegeneration through two alternative gain-of-function mechanisms. One proposed mechanism involves RNA toxicity through formation of pathogenic repeat RNA foci and sequestration of RNA-binding proteins (11-15). The other proposed mechanism for neurotoxicity of the repeats is via generation of dipeptide repeat proteins (DPRs) produced by aberrant repeat associated non-ATG (RAN) translation (16-18). Loss-of-function of the *C9orf72* gene, which encodes a guanine nucleotide exchange factor implicated in autophagy and immune responses, exacerbates gain-of-function toxicity (19-24). Work in model organisms has strongly supported the notion that DPRs play a dominant role in the pathogenesis of *C9orf72* disease, but other recent studies suggest that pathogenesis may be the result of multiple non-mutually exclusive mechanisms (25-27). Indeed, both gain-of-function mechanisms, RNA toxicity and DPR production, may independently cause disruption of pathways such as stress granule formation and nucleocytoplasmic transport (27-32). Interestingly, previous work has established that Zfp106 binds directly and specifically to GGGGCC RNA repeats, and that co-expression of Zfp106 suppresses the pathology and neurotoxicity induced by expression of 30 copies of GGGGCC in a Drosophila model of *C9orf72* ALS (2).

GGGGCC sense strand RNA repeats are capable of forming G-quadruplex or RNA hairpin structures (12, 33, 34), while antisense GGCCCC repeats form i-motifs and protonated hairpins (35). Stem-loops or hairpins are the most common RNA secondary structures and play important roles in transcription, RNA processing, mRNA export, mRNA stability and translation (36). G-quadruplexes are secondary structures that form specifically in guanine (G)-rich nucleic acids, where a tetrad of G-rich strands is coordinated by a cation (37). These secondary structures in RNA have been proposed to play multiple diverse and important roles in RNA biology, including roles in splicing, polyadenylation, transcriptional termination, translational enhancement or repression, and mRNA trafficking (37). In the context of *C9orf72* ALS/FTD, GGGGCC repeat-induced RNA foci form by phase separation, which has been suggested to be dependent on the G-quadruplex structure (30, 38, 39). Interestingly, small molecules that bind to hairpin or G-quadruplex structures formed by GGGGCC repeats reduce both DPR translation and formation of foci, demonstrating that targeting secondary structures formed by GGGGCC repeats may be therapeutically relevant (33, 40, 41).

Here, we show that Zfp106 binds to G-quadruplex-forming RNA sequences, including to GGGGCC repeats when folded in the G-quadruplex conformation. Moreover, we find that Zfp106 binding alters the G-quadruplex structure of GGGGCC repeats, and we show that Zfp106 specifically inhibits formation of GGGGCC-mediated RNA foci and suppresses GGGGCC-induced RAN translation in mammalian cells. These results support a role for Zfp106 in suppressing both of the gain-of function mechanisms associated with *C9orf72* ALS/FTD and suggest that Zfp106 recognition and binding to G-quadruplexes may play a role in the pathobiology of neurodegeneration, including *C9orf72* ALS/FTD.

## RESULTS AND DISCUSSION

### Zfp106 suppresses GGGGCC-induced RAN translation

To gain insight into how Zfp106 suppresses the cytopathic effects induced by expression of GGGGCC repeats, we co-transfected a Zfp106 expression plasmid with a pCMV-(GGGGCC)_30_-EGFP expression plasmid or a control pCMV plasmid with only 3 copies of the GGGGCC repeat sequence into Neuro-2A cells and measured eGFP RNA and protein expression in the presence or absence of Zfp106 expression (Fig. 1A). Zfp106 significantly inhibited protein expression from the eGFP mRNA containing 30 copies of the GGGGCC repeats without affecting the steady state level of the mRNA (Fig. 1A’,A’’). Zfp106 did not affect eGFP protein expression from plasmids containing 6× or 60× Huntington’s disease-associated CAG repeats (42, 43) in the 5’-UTR of the eGFP cDNA (Fig. 1B). These data suggest that Zfp106 inhibited translation of the eGFP mRNA specifically containing 30× GGGGCC repeats.

**Figure 1.**
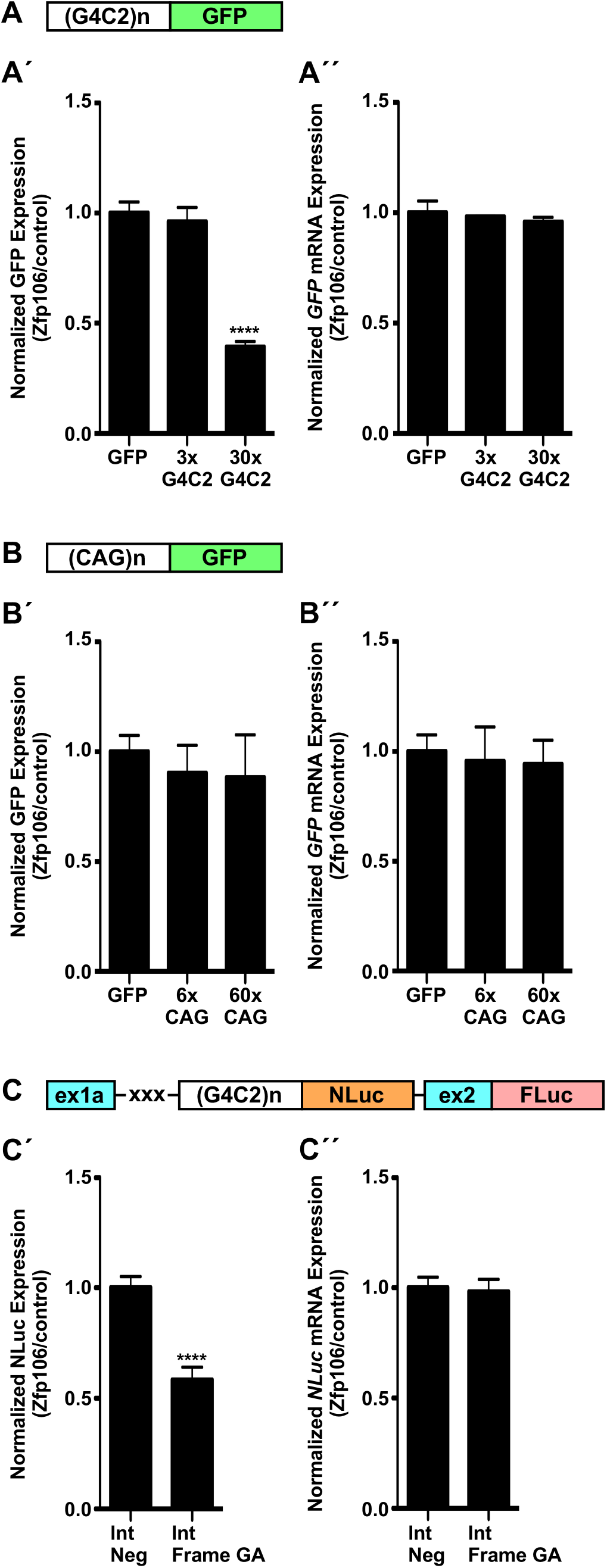
Zfp106 suppresses RAN translation. eGFP fluorescence (A’,B’) or NLuc luminescence (C’) and mRNA levels (A’’,B’’,C’’) in Neuro-2a cells (A,B) or HEK293T cells (C) co-transfected with pCMV-(GGGGCC)_n_-EGFP constructs (no-repeats, 3×, or 30× GGGGCC repeats) and Zfp106 or control vector (A); co-transfected with pCMV-(CAG)n-EGFP–containing constructs (no-repeats, 6×, or 60× CAG repeats) and Zfp106 or control vector (B); or co-transfected with bicistronic splicing dual-luciferase reporters with *C9orf72* exons and intron (Int Neg, no repeats; Int Frame GA, 70× GGGGCC repeats) and Zfp106 or control vector (C). (A,B) GFP fluorescence was normalized to total protein content; *eGFP* mRNA was normalized to *Gapdh*. (C) NLuc luminescence (C’) or mRNA expression (C’’) was normalized to FLuc in each sample. Data are expressed as ratios of Zfp106-transfected cells over control. The mean ratio from the no-repeat construct was set to a normalized value of 1. Data were analyzed by two-way ANOVA and Bonferroni’s Multiple Comparison Test; **** p<0.0001. Schematic depictions of transfected constructs are shown in (A,B,C); xxx indicates stop codons.

An important distinction between the expression of the GGGGCC repeats from the pCMV-(GGGGCC)_30_-EGFP reporter compared to the expression of the repeats in the context of *C9orf72* ALS/FTD is that the repeats expressed from the pCMV-(GGGGCC)_30_-EGFP reporter are present in the 5’-UTR, and GFP protein expression occurs by standard AUG-dependent translation (44), whereas translation of the repeats in the disease context in humans occurs via RAN translation (16-18). Therefore, to determine if Zfp106 co-expression also inhibits RAN translation, we co-transfected the Zfp106 expression plasmid with a bicistronic reporter plasmid that produces NanoLuc luciferase (NLuc) by RAN translation induced by 70× GGGGCC repeats from an intronic sequence derived from the human *C9orf72* gene (Fig. 1C, Fig. S1A) (45). As an internal control, the reporter construct produces firefly luciferase (FLuc) by canonical AUG-dependent initiation (45). Co-expression of Zfp106 significantly and specifically reduced RAN translation of NLuc from the reporter construct without affecting AUG-dependent translation of FLuc from the same plasmid (Fig. 1C’). Inhibition occurred in a Zfp106 dose-dependent manner (Fig. S1A’). Steady-state NLuc mRNA levels were unaffected by co-expression of Zfp106 (Fig. 1C’’). Zfp106 also inhibited RAN translation from a second bicistronic reporter plasmid from which NLuc is produced by RAN translation induced by 70× GGGGCC repeats from a mature mRNA, rather than from intronic sequences, also without affecting steady-state RNA level (Fig. S1B). Importantly, these data demonstrate that Zfp106 suppresses translation, including RAN translation, of GGGGCC-repeat-containing RNAs. Moreover, since DPRs produced via GGGGCC-repeat-induced RAN translation are major contributors to *C9orf72* pathology in animal models and are strongly associated with disease in humans (16-18), these results suggest that Zfp106 inhibition of RAN translation may contribute to the neuroprotective effect of Zfp106 *in vivo*.

### Zfp106 inhibits the formation of GGGGCC-repeat-induced RNA foci

GGGGCC-repeat-containing RNAs also form foci in affected motor neurons, and these foci are strongly associated with pathology and disease (11-15). Moreover, the presence of RNA foci and DPRs are not mutually exclusive, and the two pathology-associated phenomena often co-occur (46). Therefore, we examined the influence of Zfp106 co-expression on the formation of RNA foci induced by overexpression of GGGGCC-repeat-containing RNAs (Fig. 2). Jain and Vale (2017) described a system in which 29× GGGGCC repeats tagged with 12× MS2-hairpin loops can be visualized via co-expression of YFP-tagged MS2-coat binding protein, which binds to the MS2 hairpin loops (Fig. 2A) (38). Consistent with previously published results, induction of 29× GGGGCC RNA repeats in this system resulted in formation of numerous nuclear foci (Fig. 2B) (38). Remarkably, the presence of foci was substantially reduced by co-expression of Zfp106 compared to nuclear-RFP control transfected cells (Fig. 2C,D,F). In contrast, Zfp106 had no effect on the formation of RNA foci induced by 47× CAG repeats (Fig. 2E,G). As an alternate method for detecting foci, we also examined the number of foci per nucleus by using RNA FISH probes directed against MS2 hairpin loops, which also showed that Zfp106 expression substantially abrogated formation of foci when compared to the number of foci in cells lacking Zfp106 expression (Fig. S2).

**Figure 2.**
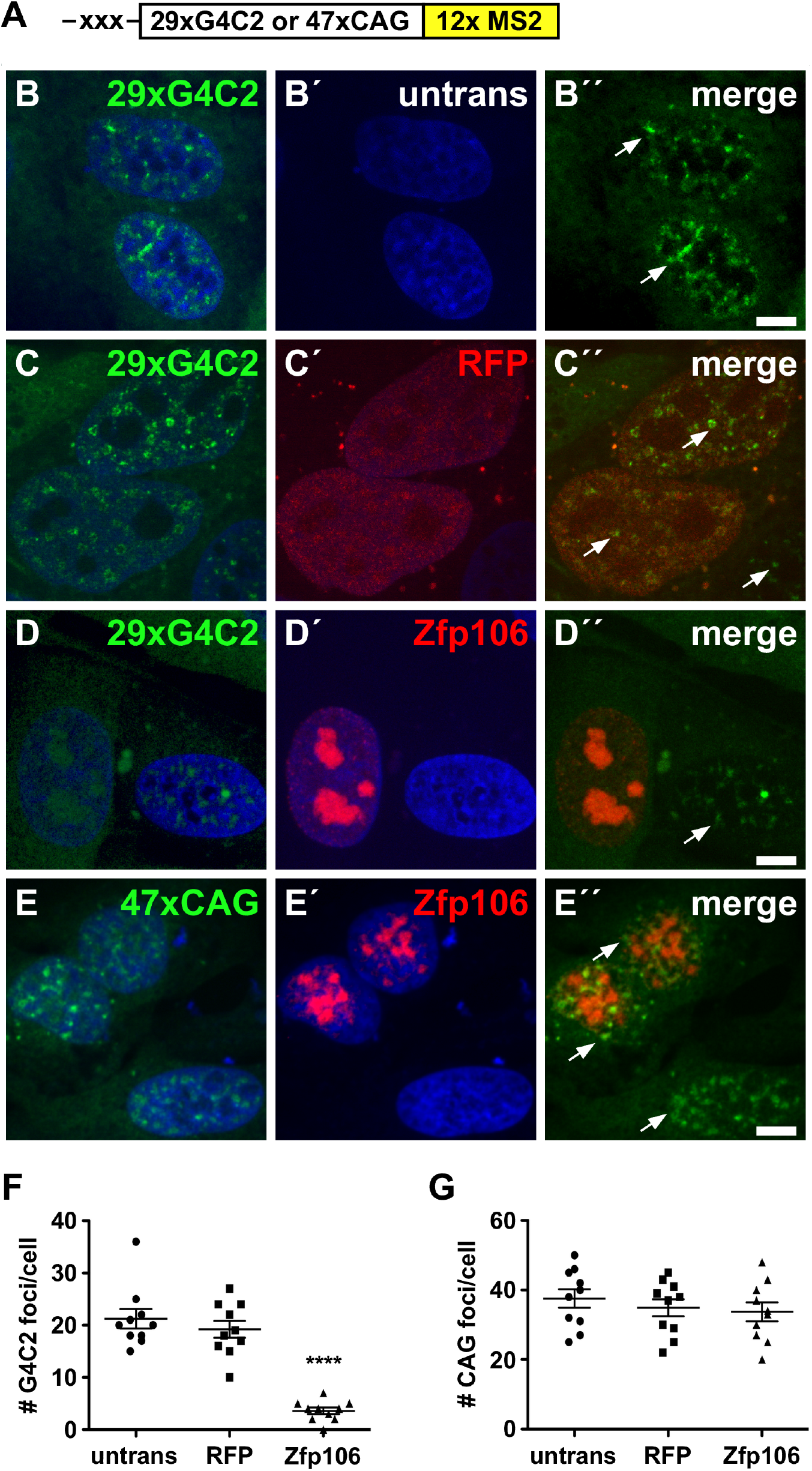
Zfp106 suppresses formation of GGGGCC-containing RNA foci. Representative fluorescence images of U-2OS cells, stably expressing either 29× GGGGCC (B,C,D) or 47× CAG repeats (E) fused to 12× MS2 RNA stem-loops (schematic shown in A), that are untransfected (B), or have been transfected with nuclear RFP (C) or RFP-Zfp106 (D,E). Cell nuclei were counterstained with DAPI (blue). (B-E) green and blue channel merge; (B’-E’) red and blue channel merge; (B’’-E’’) green and red channel merge. Arrows mark cells with >10 GGGGCC nuclear foci. Scale bars, 10 μm. Quantification of GGGGCC RNA repeat foci (F) and CAG RNA repeat foci (G) per cell analyzed by one-way ANOVA and Bonferroni’s Multiple Comparison Test. Zfp106 co-expression significantly reduced the number of GGGGCC RNA foci (F) but had no significant effect on the formation of CAG RNA foci (G); **** p<0.0001.

Taken together, the results in Figs. 1 and 2 demonstrate that Zfp106 inhibits RAN translation and the formation of RNA foci caused by GGGGCC repeats. These observations have important implications for understanding the role of Zfp106 in neuroprotection and neurodegeneration in gain- and loss-of-function models of ALS.

### Zfp106 binds to G-quadruplex-forming RNA sequences

The observations that Zfp106 suppressed RAN translation and formation of RNA foci caused by GGGGCC repeats, combined with our previously published work showing that Zfp106 binds directly and specifically to GGGGCC repeats (2), suggested that RNA-binding by Zfp106 might be central to its repressive effect on repeat-induced pathology. Our previous studies also showed that Zfp106 interacts directly with a wide array of other RNA binding proteins, which suggests that it plays roles in cellular RNA processing and that its RNA-binding activity is unlikely to be restricted to only GGGGCC repeats (2). Because GGGGCC repeats are capable of forming G-quadruplex structures and because RNA G-quadruplexes are associated with phase separation and formation of RNA foci (34, 38), we reasoned that Zfp106 might bind more generally to G-quadruplex forming RNA sequences and that this ability might confer its repressive effects on GGGGCC-repeat toxicity.

We use electrophoretic mobility shift assay (EMSA) to test the ability of Zfp106 to bind to numerous RNA repeat sequences, including those predicted to form G-quadruplexes and those predicted to form hairpins or other non-G-quadruplex secondary structures (Fig. 3A). Zfp106 bound to all sequences predicted to form G-quadruplexes, while it failed to bind significantly to all non-G-quadruplex-forming sequences (Fig. 3A). Indeed, Zfp106 bound efficiently to GGGGCC repeats under conditions that promote G-quadruplex formation (Fig. 3A). In contrast, under those same conditions, Zfp106 showed no binding to antisense GGCCCC repeats (non-G-quadruplex-forming), CUG repeats, GGGGCC repeats with G>A substitutions that impede G-quadruplex formation (MUT), or AAAACC repeats (Fig. 3A). Likewise, Zfp106 showed little to no detectable binding to a GC stem-loop sequence (Fig. 3A’’) or to CAG repeats (Fig. 3B), consistent with the observations that Zfp106 did not suppress translation or formation of foci from transcripts containing CAG repeats (Fig. 1B, Fig. 2E,G).

**Figure 3.**
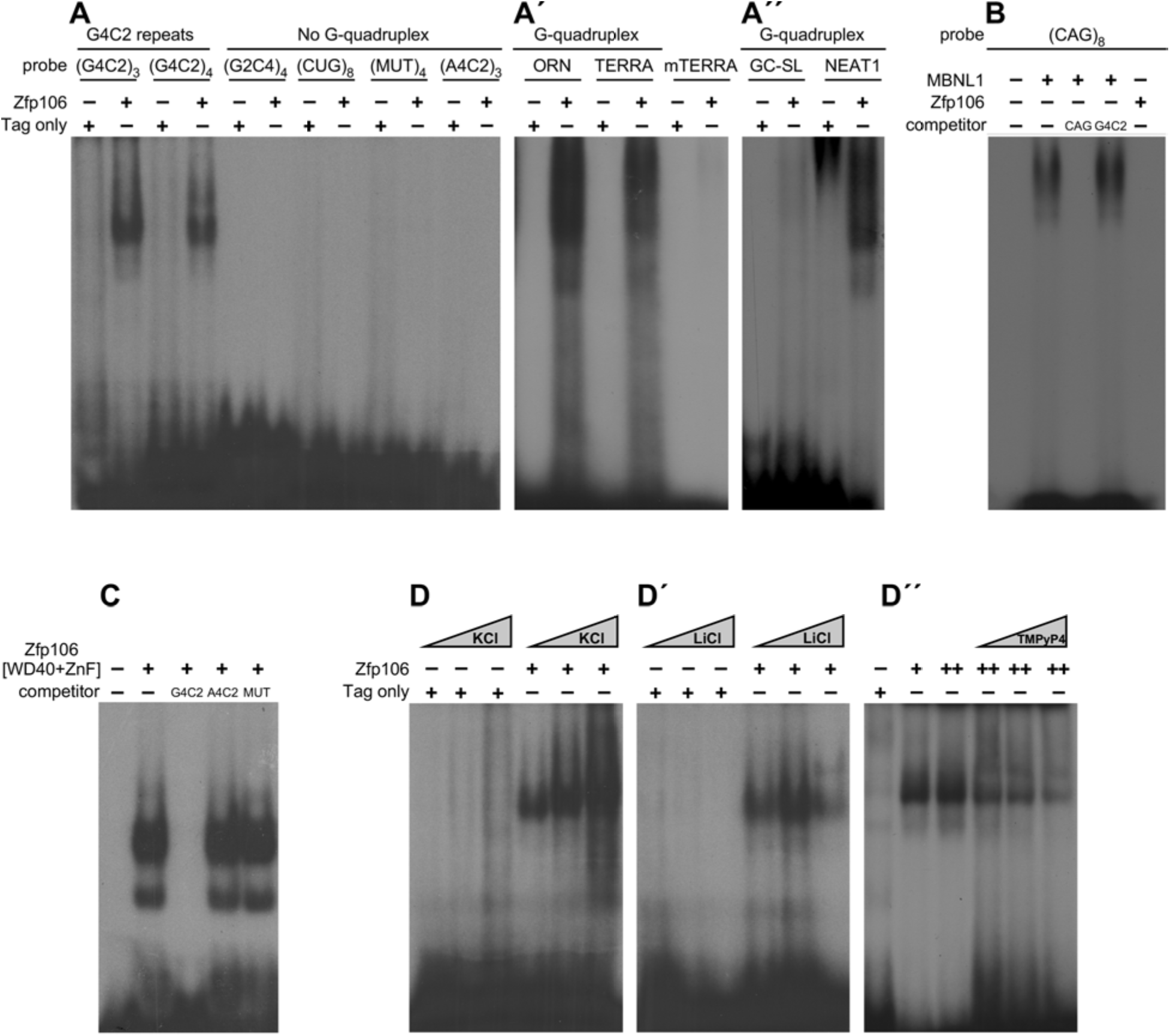
Zfp106 binds G-quadruplex RNAs. (A-A’’) RNA EMSAs performed with purified Zfp106 protein demonstrate that Zfp106 directly binds G-quadruplex forming RNAs in *vitro*: GGGGCC [(G4C2)_3_, (G4C2)_4_] repeats (A), ORN and TERRA (A’) and NEAT1 (A’’). Zfp106 showed no binding to the following non-G-quadruplex-forming RNAs: antisense GGCCCC repeats (G2C4)_4_, (CUG)_8,_ mutated GGGGCC [(MUT)_4_], (A4C2)_3_, mutated TERRA (mTERRA), and GC-rich stem-loop (GC-SL). (B) RNA EMSA using purified Zfp106 protein demonstrates that Zfp106 does not bind (CAG)_8_ repeats under conditions in which the RNA binding protein MBNL1 efficiently binds (CAG)_8_. MBNL1 binding to (CAG)_8_ is specific since binding is competed by 30× molar excess of self-competitor but not 30× molar excess of (GGGGCC)_4_. (C) RNA EMSA using purified Zfp106 C-terminus [WD40+ZnF] demonstrates that [WD40+ZnF] is sufficient to bind to (GGGGCC)_4_ repeats in vitro. 30× molar excess of unlabelled self-competitor, but not 30× molar excess of unlabeled (AAAACC)_4_ or (MUT)_4_, efficiently competed with [WD40+ZnF] binding to (GGGGCC)_4_, establishing specificity of the interaction. (D-D’’) RNA EMSA using purified Zfp106 protein shows increased binding of Zfp106 to (GGGGCC)_4_ with increasing concentration of KCl (3 – 30 – 100 mM) (C) and decreased binding with increasing concentrations of LiCl (3 – 30 – 100 mM) and TMPyP4 (10 – 25 – 50µM).

The full length isoform of Zfp106 used in these studies is ∼150kD, comprises 1,099 amino acids, and is composed primarily of an N-terminal zinc finger, predicted low complexity domains, seven WD-40 domains and two C-terminal zinc fingers (Fig S3A) (3, 47). To gain additional insight into RNA binding by Zfp106, we next sought to identify determinants in Zfp106 responsible for binding to GGGGCC repeats. We found that a C-terminal 43kD-fragment of Zfp106, comprising amino acid residues 712-1099 and encompassing the WD40 domains and the two C-terminal zinc fingers was sufficient for robust, specific binding to (GGGGCC)_4_ repeats (Fig. 3C). Other regions of Zfp106, including a fragment encoding the N-terminus and a fragment including N-terminal and C-terminal zinc fingers, but lacking intervening low complexity regions, showed no detectable binding to (GGGGCC)_4_ repeats (Fig. S3A,B).

GGGGCC RNA repeats form parallel G-quadruplexes (12). In addition to binding to (GGGGCC)_4_ and (GGGGCC)_3_ repeats, Zfp106 also bound efficiently to other parallel G-quadruplex-forming RNA sequences, including sequences in TERRA and NEAT1 (Fig. 3A’,A’’) (48, 49). Mutation of TERRA repeats with G>U substitutions, which impede G-quadruplex formation, prevented Zfp106 binding (Fig. 3A’). Interestingly, the predicted anti-parallel G-quadruplex-forming sequence from ORN-N (50) was also bound by Zfp106 (Fig. 3A’). Zfp106 binding to NEAT1 RNA is particularly interesting since GGGGCC RNA repeats have been suggested to replace NEAT1 RNA as a scaffold in paraspeckle-like structures and to form foci with characteristics of paraspeckles (51, 52). Moreover, the DPR protein poly-PR binds to and up-regulates NEAT1, resulting in increased paraspeckle formation, which might contribute to the neurotoxic effect of poly-PR (53).

As further evidence for the specificity of Zfp106 binding to G-quadruplex-forming structures, rather than to primary sequences in RNA, we found that addition of KCl to Zfp106-(GGGGCC)_4_ EMSA increased Zfp106 binding, whereas addition of LiCl to Zfp106-(GGGGCC)_4_ EMSA decreased Zfp106 binding (Fig. 3D,D’). This is significant because it is well understood that G-quadruplexes are differentially stabilized or destabilized by monovalent cations, with potassium having a stabilizing effect and lithium exerting a destabilizing effect (54, 55). Similarly, addition of increasing concentration of TMPyP4, a cationic porphyrin derivative that binds and distorts the G-quadruplex structure of GGGGCC repeats (56, 57), decreased Zfp106 binding to (GGGGCC)_4_ repeats (Fig. 3D’’).

### Zfp106 alters the G-quadruplex structure of GGGGCC repeats upon RNA binding

The threshold number of ∼30 GGGGCC repeats for disease is similar to the repeat length in which cells in culture exhibit foci, but higher than the number of repeats required for RNA gelation *in vitro* (38). For instance, as few as 5× GGGGCC RNA repeats form spherical, gel-like clusters *in vitro*, but this number of repeats is clearly not sufficient for formation of foci or phase separation *in vivo* (38). The discrepancy between the number of repeats required for phase transition and gelation *in vitro* versus *in vivo* may be due to cellular proteins that alter the G-quadruplex structure in a way that prevents phase separation *in vivo* (38, 58). Therefore, we reasoned that the disruption of RNA foci and suppression of neurotoxicity caused by GGGGCC repeats in cells and flies might be due to the ability of Zfp106 to bind to and alter the RNA G-quadruplex secondary structure. To test this idea, we analyzed the interaction of the C-terminal 43kD-fragment of Zfp106 (Zfp106[WD40+ZnF]) with GGGGCC repeats by circular dichroism (CD). CD spectroscopy confirmed that the (GGGGCC)_4_ RNA molecules formed the expected parallel G-quadruplex structure, with a spectral minimum at 236 nm and a spectral maximum at 264 nm (Fig. 4A). The CD spectra for Zfp106[WD40+ZnF] alone showed negative peaks in the 210-220nm range but gave minimal to no signal at 264 nm (Fig. 4C), allowing us to monitor the G-quadruplex structure of (GGGGCC)_4_ in the presence and absence of Zfp106 during temperature unfolding (Fig. 4A,B).

**Figure 4.**
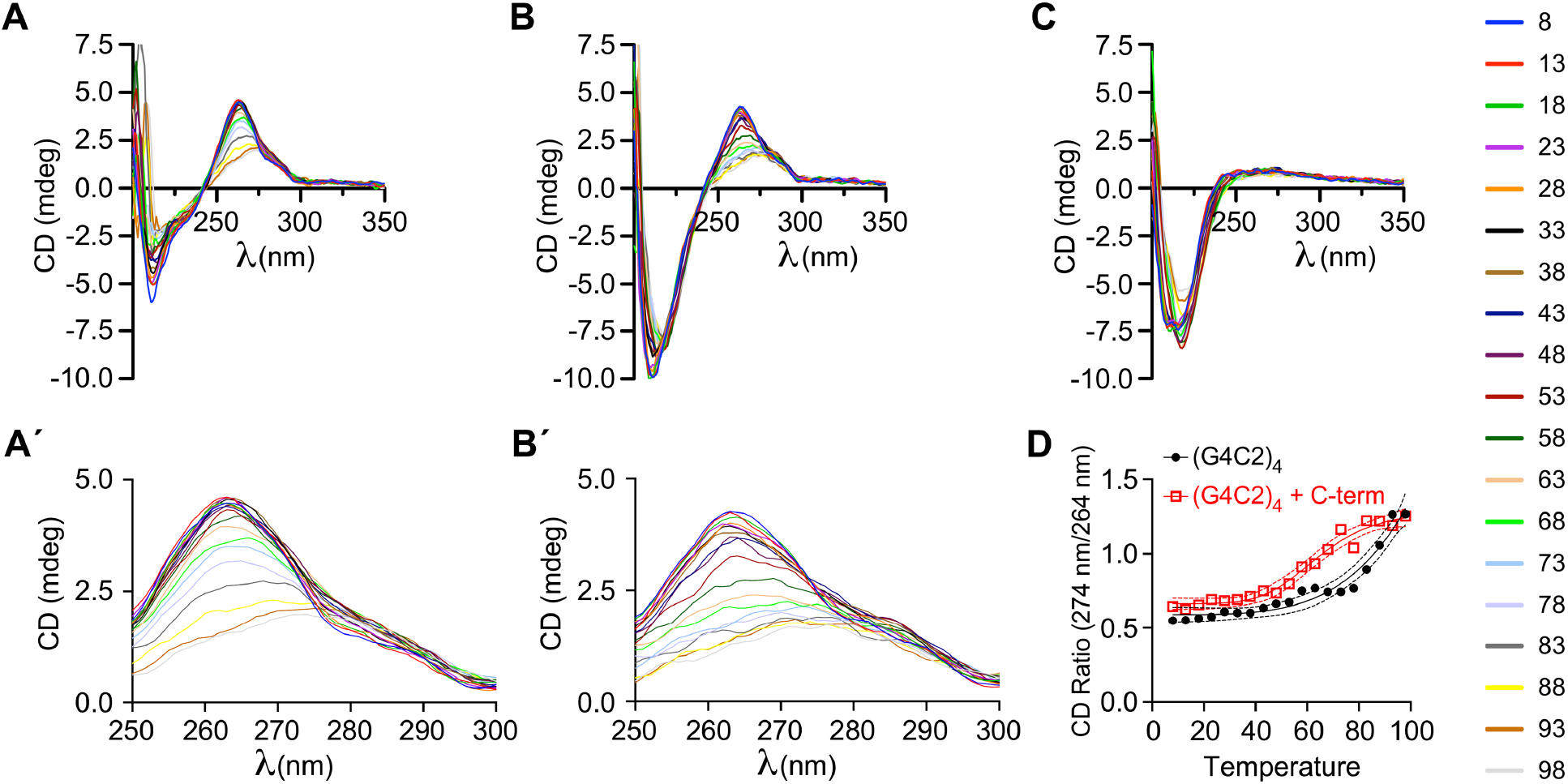
Zfp106 [WD40+ZnF] alters the G-quadruplex structure of r(GGGGCC)_4_. (A,A’) CD spectra for (GGGGCC)_4_ RNA showing a characteristic parallel G-quadruplex structure with a minimum at 236 nm and a maximum at 264 nm. (B,B’) CD spectra for r(GGGGCC)_4_ plus Zfp106 C-terminus [WD40+ZnF]. (C) CD spectra for Zfp106 C-terminus [WD40+ZnF] alone. Zfp106 C-terminus binding to GGGGCC)_4_ RNA repeats caused a change in shape of the CD spectra in the 250-300 nm region with a shift of the peak at 264 nm to 274 nm (A’,B’). CD spectra were measured during a thermal unfolding with increasing temperature from 8 to 98°C, depicted by the color scale on the right of the figure. (D) Ratio of CD absorbance at 274 nm/264 nm from 8 to 98°C shows that binding of Zfp106 C-terminus [WD40+ZnF] caused a conformational shift in the G-quadruplex structure with a shift from absorbance at 264 nm to 274 nm occurring at lower temperature (red squares) than in control (black circles).

Addition of the C-terminal 43kD-fragment of Zfp106 (Zfp106[WD40+ZnF]) at an equimolar ratio of protein to RNA resulted in a conformational change in the (GGGGCC)_4_ G-quadruplex structure, reflected by the lowering and broadening of the peak at 264 nm and by a shift and change in shape of the CD spectra in the 250-300 nm range (Fig. 4A’,B’). Closer examination of CD spectra in the 250-300 nm range showed a shift in the peak from 264 nm to 274 nm for G4C2 repeats alone as the temperature increased, suggesting a change in the G-quadruplex conformation (Fig. 4 A’). Interestingly, this shift in the peak from 264 nm to 274 nm occurred at lower temperature in the presence of Zfp106 (Fig. 4B’), suggesting that Zfp106 stabilized the G-quadruplex conformation normally favored at higher temperature. This observation was more clearly indicated by comparing the ratio of the CD absorption at 274 nm to 264 nm in the presence and absence of Zfp106 C-terminus (Fig. 4D). We also conducted essentially identical experiments with full-length Zfp106 at 1:2 molar ratio of protein to RNA, and similar results were obtained (Fig. S4), confirming a change in the (GGGGCC)_4_ G-quadruplex structure upon full-length Zfp106 binding.

The ability of Zfp106 to induce a conformational change in the G-quadruplexes formed by GGGGCC repeats is particularly interesting given prior studies that have suggested that phase transition and formation of RNA foci is dependent on the G-quadruplex structure (30, 38, 39). Indeed, as discussed earlier, small molecules targeting the G-quadruplex structure of GGGGCC repeat RNAs can reduce RNA foci formation, DPR levels and GGGGCC-induced toxicity (33, 41). It has been suggested that (GGGGCC)_4_ might form both intermolecular and intramolecular G-quadruplexes in a concentration- and temperature-dependent manner (59). However, when we analyzed (GGGGCC)_4_ repeats by analytical ultracentrifugation, we failed to detect intermolecular interactions between (GGGGCC)_4_ molecules under the same buffer conditions and RNA concentration used in our CD analysis (data not shown).

Thus, our data suggest that Zfp106 binding to GGGGCC repeats induces a shift in the intramolecular G-quadruplex structure, although we cannot rule out a possible effect on intermolecular forms at higher concentrations of GGGGCC repeats such as might occur *in vivo*. Regardless, the studies presented here support the idea that Zfp106’s suppressive effect on GGGGCC-mediated cytotoxicity might be through alteration or stabilization of specific G-quadruplex structures formed by GGGGCC repeats. Future studies will dissect how different G-quadruplex conformations contribute to disease in C9orf72 ALS.

### GGGGCC repeats may exert a “sponge”-like effect on Zfp106 or other G-quadruplex RNA-binding proteins

Previous studies have demonstrated that GGGGCC RNA repeats sequester various G-quadruplex-binding proteins involved in diverse cellular functions, and this sequestration may contribute to the cytotoxic effects of the repeats (11, 60). For example, the G-quadruplex binding protein nucleolin is sequestered by GGGGCC RNA repeats, resulting in nucleolin mislocalization and nucleolar stress in *C9orf72* ALS patients (12). Similarly, sequestration of the G-quadruplex binding protein hnRNPH/F by GGGGCC RNA repeats results in defective RNA splicing and polyadenylation (11, 13, 14). GGGGCC repeats have also been shown to compromise nucleocytoplasmic transport by sequestering the RBP RanGAP1 (61). Thus, it is interesting to speculate that part of the toxic effects of GGGGCC repeats in the context of *C9orf72* ALS/FTD may be due to sequestration of Zfp106. Indeed, loss of *Zfp106* function in mice causes severe ALS-like neuromuscular disease (1, 2, 7), supporting the idea that Zfp106 sequestration could contribute to neurodegenerative disease. However, precisely how Zfp106 loss-of-function or sequestration leads to neurodegeneration is unclear since the specific function of Zfp106 in RNA processing and metabolism remains to be determined. The observations that Zfp106 binds to G-quadruplex RNAs and induces conformational changes in the G-quadruplex structure of GGGGCC repeats suggests that Zfp106 might play a role in G-quadruplex RNA homeostasis and metabolism, as has been suggested for other G-quadruplex binding proteins (62). Future studies will address the functional importance of Zfp106 binding to G-quadruplex-forming cellular RNAs and if dysregulation of the “G4ome” (62) is a contributing mechanism for neurodegeneration in *C9orf72* ALS/FTD or other neurodegenerative diseases.

## MATERIALS AND METHODS

### Plasmids

GFP reporter constructs with either 3× or 30× GGGGCC repeats inserted in the 5’-UTR of an eGFP cDNA in plasmids pCMV-(GGGGCC)_3_-EGFP and pCMV-(GGGGCC)_30_-EGFP, respectively, have been described previously (44). Plasmid pCMV-(CAG)_6_-EGFP was generated by cloning a synthetic oligonucleotide for 6× CAG repeats into the EcoRI and KpnI sites in between the transcription site and the translation site of pEGFP-N3, as previously described for GGGGCC repeats (44). Similarly, plasmid pCMV-(CAG)_60_-EGFP was generated by cloning 30× CAG repeats into the EcoRI and KpnI sites of pEGFP-N3, followed by cloning of an additional 30× CAG repeats into the XhoI and KpnI sites of pEGFP-N3, also as previously described for GGGGCC repeats (44). Bicistronic reporters pcDNA5 FRT TO FLuc2-noC9R-NLuc (Neg), pcDNA5 FRT TO FLuc2-C9R70-NLuc Frame GA (Frame GA) and bicistronic splicing reporters pcDNA5 FRT TO noC9R intron-NLuc-exFLuc 2 (Int-Neg) and pcDNA5 FRT TO C9R70 intron-NLuc-exFLuc 2 Frame GA (Int-Frame GA) have been described previously (45).

Plasmid pCMV-GFP-Zfp106 was created by cloning the Zfp106 cDNA into the XhoI-EcoRI sites in pEGFP-C2 (Clontech). Plasmid pCMV-RFP-Zfp106 was created by replacing EGFP in plasmid pCMV-GFP-Zfp106 with NLS-TagRFP. 3×FLAG-2×STREP-Zfp106 has been described previously (2). Truncation and deletion mutants of Zfp106 were generated by PCR and confirmed by sequencing; Zfp106 cDNA fragments, encoding amino acids 712-1099 of the C-terminal region, encoding amino acids 1-711 of the N-terminal region of the full-length protein, and a cDNA with an internal deletion such that the resultant cDNA encodes amino acids 1-178 and 1025-1099 were digested and subcloned into the NotI and XhoI sites of pcDNA4/TO (Invitrogen) modified to include N-terminal 3×FLAG-2× STREP tags to create plasmids 3×FLAG-2×STREP-Zfp106[Cterm], 3×FLAG-2×STREP-Zfp106[Nterm], and 3×FLAG-2×STREP-Zfp106[N+Cterms], respectively. 3×FLAG-2×STREP-Zfp106[Cterm] encodes Zfp106[WD40+ZnF].

### Cell culture, transfections, and reporter assays

Mouse Neuro-2a (ATCC, CCL-131; RRID:CVCL_0470) and human HEK293T cells (ATCC, CRL-3216; RRID:CVCL_0063) were obtained directly from ATCC and maintained in Dulbecco’s modified Eagle Medium supplemented with 10% fetal bovine serum (FBS, Gibco, Waltham, MA) and penicillin-streptomycin (Gibco). Monoclonal U-2OS cell lines stably expressing Tet-On 3G transactivator protein (Clontech), a tandem-dimeric MS2 hairpin binding protein tagged with eYFP (MS2CP-YFP), and 29× GGGGCC or 47× CAG repeat under control of a doxycycline-inducible promoter have been described previously (38). All cell lines were routinely tested for Mycoplasma contamination with the MycoAlert PLUS Mycoplasma Detection Kit (Lonza, Switzerland, LT07-705) and found to be negative.

Neuro-2a and U-2OS cells were transfected using GenJet In Vitro DNA Transfection Reagent for Neuro-2A Cells or U-2OS respectively (Signagen); HEK293T cells were transfected using PolyJet DNA In Vitro Transfection Reagent (Signagen), according to the manufacturers’ recommendations. Protein purification from HEK293T cells was performed as described previously (2).

For reporter assays, three biological replicates were analyzed for each condition for all GFP and dual-luciferase experiments. For bicistronic reporters, gene expression was induced with 2 μg/ml doxycycline for 24 h prior to sample collection. Nano-luciferase (NLuc) and firefly luciferase (FLuc) activities were measured by using the Nano-Glo Dual Luciferase Assay on a Glomax Multidetection System (Promega). GFP fluorescence was measured using an Infinite M1000 microplate reader (Tecan Group Ltd); samples were normalized by total protein concentration using the Bradford assay with protein assay dye reagent (Bio-Rad).

### qPCR, imaging of RNA foci, and fluorescence in situ hybridization (FISH)

For reverse transcriptase (RT)-qPCR, total RNA was isolated from cells using Trizol (Invitrogen) and was treated for 1 h at 37°C with DNaseI, followed by cDNA synthesis using random primers and the Omniscript RT kit (Qiagen). qPCR was performed using the MAXIMA SYBR Green kit (Thermo Scientific) and a 7900HT Fast Real Time PCR System or a Quant Studio 5 Real Time PCR System (Applied Biosystems) with the primers listed in Supplemental Table S1. Data were normalized to housekeeping gene *Gapdh* by the 2−ΔΔCt method (63).

Repeat RNA expression in U-2OS cells stably transfected with 29× GGGGCC or 47× CAG repeats was induced by adding 1μg/ml doxycycline at the time of transfection with the RFP-Zfp106 plasmid described above. 24 h post-transfection, cells were fixed in 4% paraformaldehyde for 10 minutes and imaged in 0.3 µm Z-stacks using a Nikon Ti-Eclipse microscope equipped with Yokogawa confocal CSU-X spinning disk module. The number of foci per cell was calculated using the FIJI 3D Objects Counter plugin as previously described (38).

For RNA FISH, repeat-expressing U-2OS cells were induced for 24 h and were then harvested and fixed in 2% paraformaldehyde for 10 minutes at room temperature. Following fixation, cells were permeabilized by overnight incubation in 70% ethanol at 4°C. RNA was detected using Cy5-labeled DNA oligonucleotides designed against the MS2-hairpin sequence; probe sequences were previously described (38). Stellaris RNA FISH hybridization and wash buffers were purchased from Biosearch Technologies and used according to the manufacturer’s recommendations. After labeling, samples were mounted in Prolong Gold antifade medium (Thermo Scientific, Inc) and imaged using confocal microscopy as described above.

### Electrophoretic mobility shift assay (EMSA)

100 pmol of single-stranded RNA (Integrated DNA Technologies, Inc.) were 5′-end labeled with [γ-^32^P]ATP, unincorporated radioactivity was removed, and RNA EMSA was performed as described previously (2). Full-length Zfp106 and indicated truncation and deletion mutants or MBNL1 protein were purified from total cellular lysates of transfected HEK293T using the Strep-Tag system (64) with Strep-Tactin Sepharose (IBA, catalog #2-1201-010), according to the manufacturer’s protocol; 0.1% Triton X-100 was added to all buffers for cell lysis and chromatography. Equivalent amount of total protein was used for each EMSA. Parallel purification from HEK293T cells transfected with the 3×FLAG-2×STREP parental vector was performed and the corresponding fractions in which Zfp106 and MBNL1 proteins were eluted were used in RNA EMSA as a negative control. RNA-protein mixtures were electrophoresed in 6% acrylamide-TBE gels, which were subsequently dried and subjected to autoradiography. For competition experiments, the purified proteins were preincubated for 10 minutes at room temperature with 30-fold molar excess of unlabeled single-stranded RNA competitor. The single-stranded RNA probe sequences are provided in Supplemental Table S2.

### Circular dichroism (CD) spectroscopy

CD spectra from 400 to 190 nm were collected at temperatures between 8°C and 98°C, with a 5°C/minute temperature gradient, using a Jasco J-810 equipped with a Jasco Peltier temperature control system. CD spectra were recorded in the presence of 4 μM (GGGGCC)_4_ RNA, either alone or with addition of 4 μM Zfp106 C-terminal fragment (Zfp106[WD40+ZnF]) or 2 μM full length Zfp106 in 10 mM sodium cacodylate (pH 7.35). Prior to use in CD, Zfp106 and Zfp106[WD40+ZnF] proteins were purified as described above and then buffer exchanged to 10 mM sodium cacodylate (pH 7.35) and concentrated using the Amicon Pro Purification System (Millipore, Sigma). Three scans were recorded and averaged for each sample. A CD spectrum of the buffer was recorded and subtracted from all spectra.

### Statistical analyses

Statistical analyses were performed using GraphPad Prism 5.0 (GraphPad Software; RRID:SCR_002798). Data were analyzed by one-way or two-way ANOVA followed by Bonferroni’s Multiple Comparison Test or Dunnett’s Multiple Comparison Test.

## ACKNOWLEDGEMENTS

HTK was supported by NIH grant K99GM138753. BC received support from the UCSF Program for Breakthrough Biomedical Research, which is partially funded by the Sandler Foundation. This work was supported primarily by grant AL210129 from the Department of Defense CDMRP to BLB with additional support from NIH grants R35 GM122603 to WFD and DK119621 and HL146366 to BLB. We are grateful to Peng Jin (Emory) and Shuying Sun (Johns Hopkins) for providing plasmids and Ron Vale (Janelia) and Ankur Jain (MIT) for sharing cells and for helpful advice. We thank Bill Seeley, Sarat Vatsavayai, Jen Yokoyama, and members of the Black and DeGrado labs for helpful discussions.

## AUTHOR CONTRIBUTIONS

Conceptualization, BC, WFD, and BLB; Methodology, Formal Analysis, and Investigation, BC, HW, BD, HTK; Writing – Original Draft, BC and BLB; Writing – Reviewing and Editing, all authors; Visualization, BC and BLB; Project Administration, BC and BLB; Supervision, BLB and WFD; Funding Acquisition, BLB and WFD. All authors approved the final manuscript. The authors declare no competing interests.

