## Supplemental Figures and Tables for "Zfp106 binds to G-quadruplex RNAs and inhibits RAN translation and formation of RNA foci caused by G4C2 repeats"

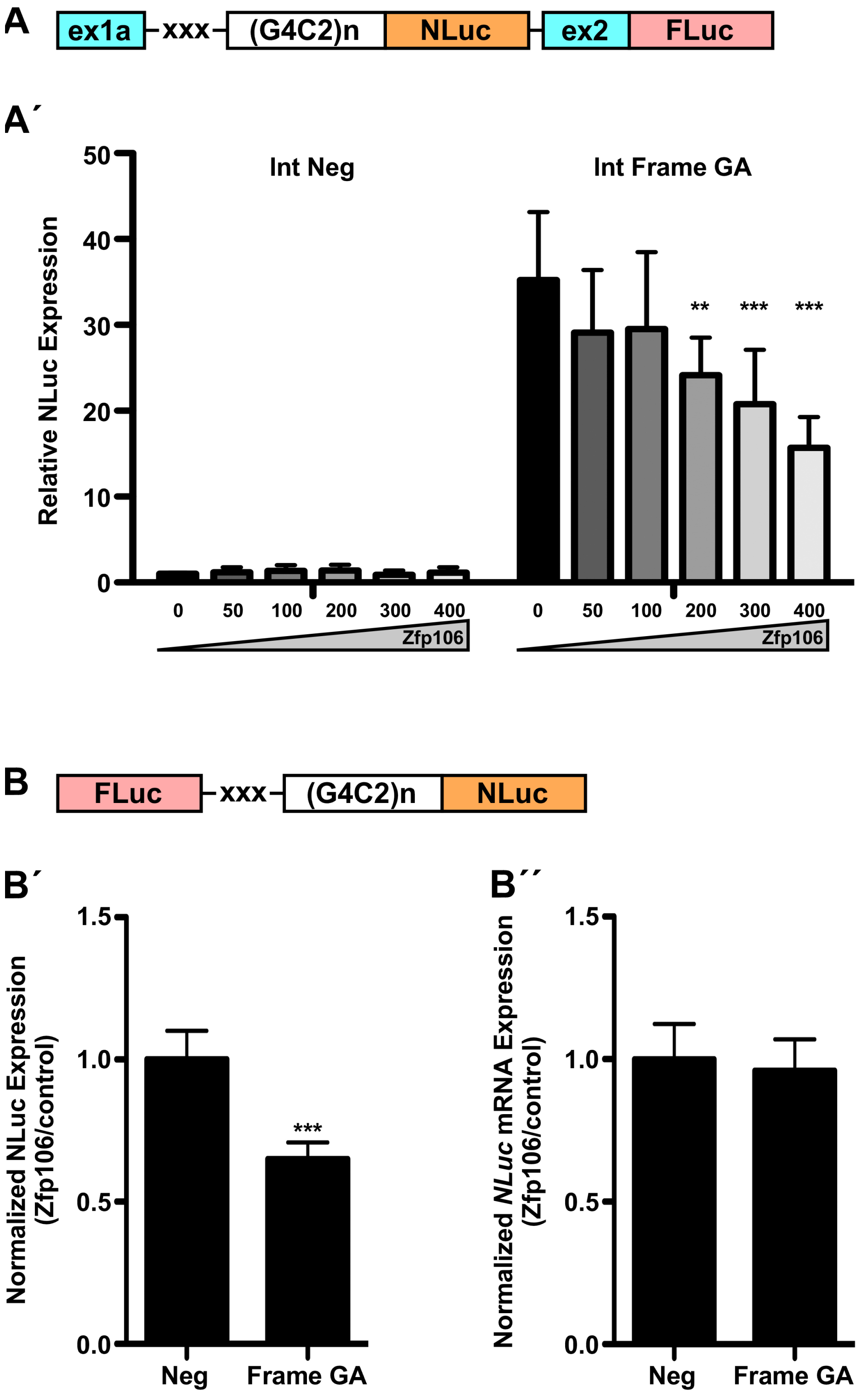
A
ex1a
xxx
(G4C2)n
NLuc
ex2
FLuc
A´
Relative NLuc Expression
Int Neg
Int Frame GA
**
***
***
0
50
100
200
300
400
Zfp106
B
FLuc
xxx
(G4C2)n
NLuc
B´
Normalized NLuc Expression (Zfp106/control)
***
Neg
Frame GA
B´´
Normalized NLuc mRNA Expression (Zfp106/control)
Neg
Frame GA

**Fig. S1. Zfp106 suppresses RAN translation.** (A) NLuc fluorescence in HEK293T cells co-transfected with bicistronic splicing dual-luciferase reporters with *C9orf72* exons and intron (Int Neg, no repeats; Int Frame GA, 70× GGGGCC repeats) and increasing amounts of Zfp106 (shown at the bottom of the graph in nanograms). Parental control vector DNA was added to each transfection to ensure an equivalent amount of plasmid DNA. No-repeat control transfected with parental control was set to a normalized value of 1. Zfp106 co-expression significantly suppressed RAN translation from the Int Frame GA construct containing 70× repeats in a dosage-dependent fashion. (B) NLuc luminescence (B′) as a readout of RAN translation and mRNA levels (B′′) in HEK293T cells co-transfected with bicistronic (no splicing) dual-luciferase reporters (Neg, no repeats; Frame GA, 70x GGGGCC repeats) and Zfp106 or control vector. NLuc luminescence was normalized to FLuc in each sample. Data are expressed as ratios of Zfp106-transfected cells over control-transfected cells. The mean ratio from the no-repeat construct (Neg) was set to a normalized value of 1. Data were analyzed by one-way ANOVA and Bonferroni's Multiple Comparison Test; ** $p<0.01$, *** $p<0.001$. Schematic depictions of transfected constructs are shown in (A,B); xxx indicates stop codons.

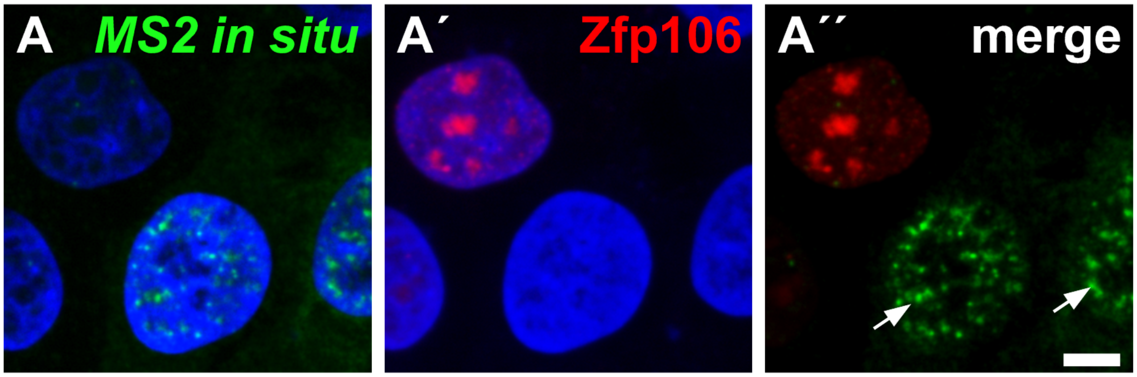


**Fig. S2. Zfp106 suppresses formation of GGGGCC-containing RNA foci.** RNA fluorescence *in situ* hybridization (FISH) using probes directed against MS2 hairpin loops confirms that Zfp106 co-expression significantly reduced the number of GGGGCC RNA foci in the 29× GGGGCC U-2OS cell line. Cell nuclei were counterstained with DAPI (blue). (A) green and blue channel merge; (A′) red and blue channel merge; (A′′) green and red channel merge. Arrows mark cells with >10 GGGGCC nuclear foci. Scale bars, 10 μm.

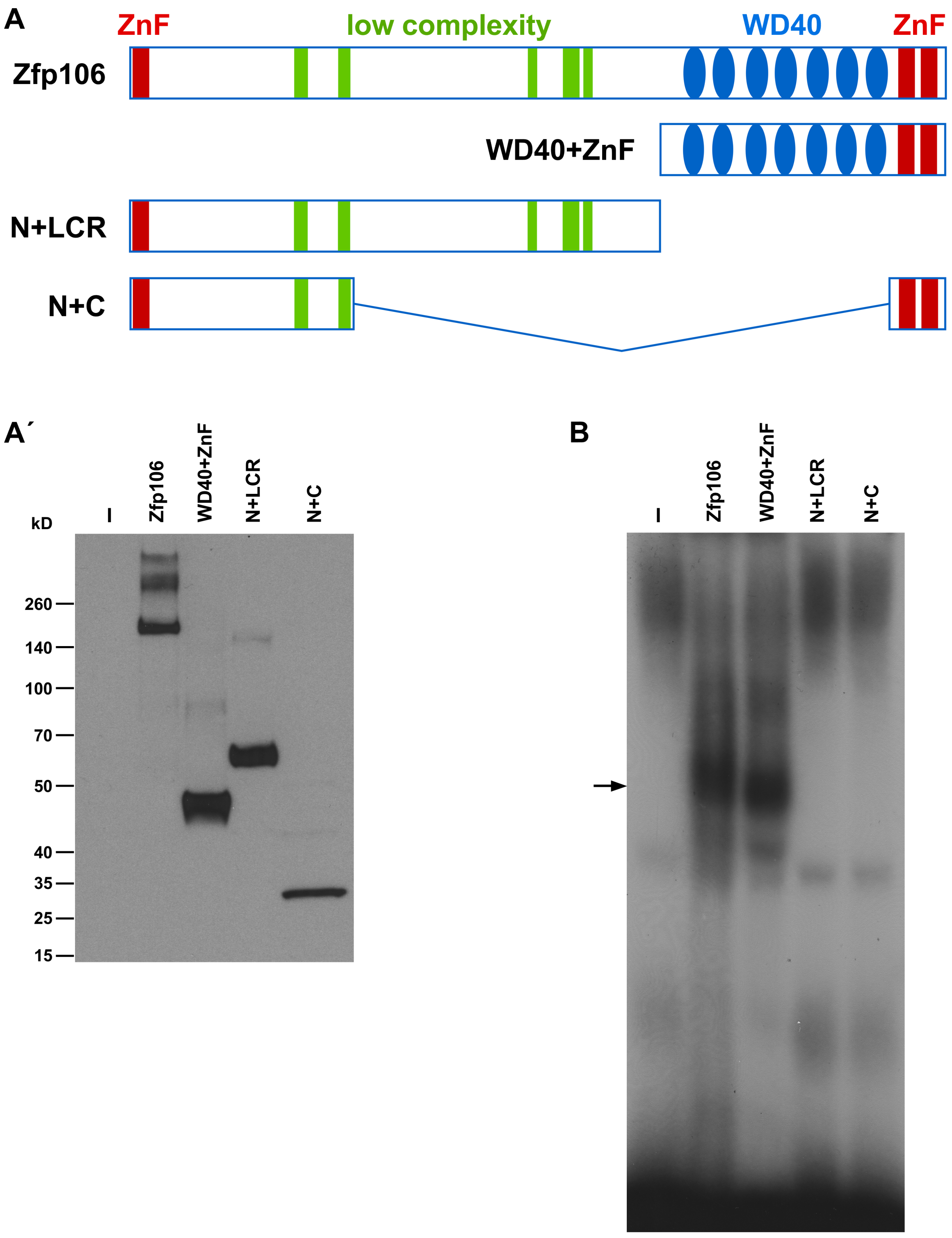


**Fig. S3. The Zfp106 C-terminus [WD40+ZnF] is sufficient for bind to r(GGGGCC)$_4$.** Schematic (A) and western blot (A′) of Zfp106 deletion constructs tested for binding to (GGGGCC)$_4$ via RNA EMSA (B). Full-length Zfp106 and [WD40+ZnF] comparably bound to r(GGGGCC)$_4$, while [the N-terminal and low complexity regions (LCR) [N+LCR] and [N+C] showed no detectable binding to r(GGGGCC)$_4$ by EMSA.

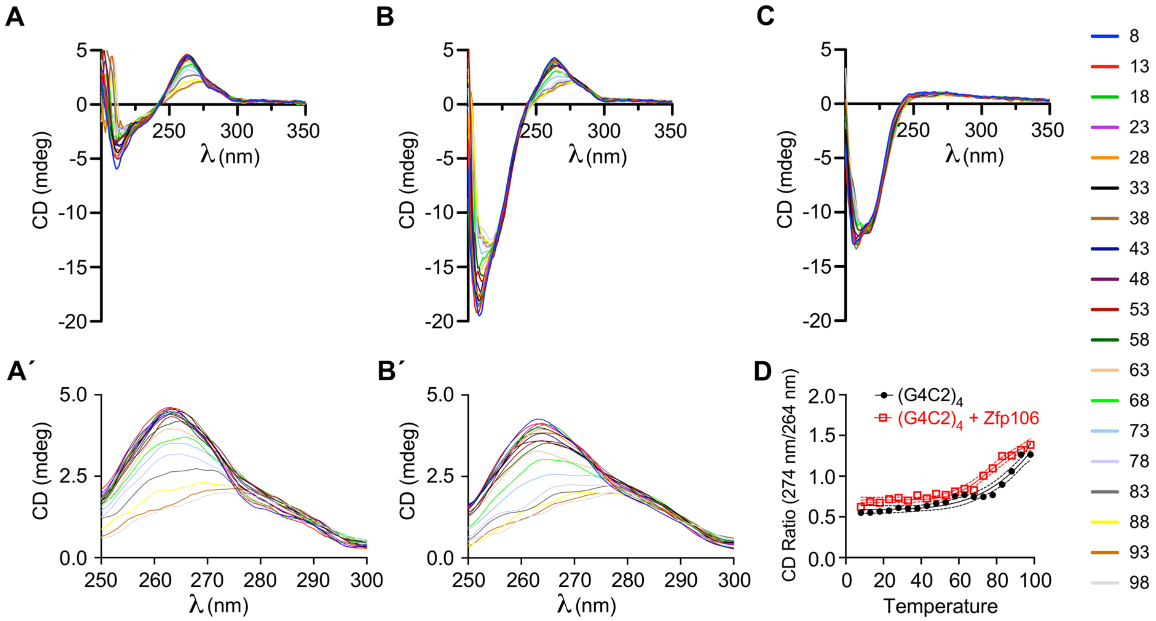


**Fig. S4. Zfp106 alters the G-quadruplex structure of r(GGGGCC)$_4$.** (A,A′) CD spectra for (GGGGCC)$_4$ RNA showing a characteristic parallel G-quadruplex structure with a minimum at 236 nm and a maximum at 264 nm. (B,B′) CD spectra for r(GGGGCC)$_4$ plus full length Zfp106. (C) CD spectra for Zfp106 alone. Zfp106 binding to GGGGCC)$_4$ RNA repeats caused a change in shape of the CD spectra in the 250-300 nm region with a shift of the peak at 264 nm to 274 nm (A′,B′). CD spectra were measured during a thermal unfolding with increasing temperature from 8 to 98°C, depicted by the color scale on the right of the figure. D) Ratio of CD absorbance at 274 nm/264 nm from 8 to 98°C shows that binding of Zfp106 caused a conformational shift in the G-quadruplex structure with a shift from absorbance at 264 nm to 274 nm occurring at lower temperature (red squares) than in control (black circles) even when present in a 1:2 molar ratio. Note: panels A,A′ are identical to panels A,A′ in Fig. 4.

Table S1. T Primers used for RT-qPCR (qPCR).

| **gene** | **5′ primer** | **3′ primer** |
|---|---|---|
| *egfp* | 5′-gcacgacttcttcaagtccgccatgcc-3′ | 5′-gcggatcttgaagttcaccttgatgcc-3′ |
| *GAPDH* (human) | 5′-gagtcaacggatttggtcgt-3′ | 5′-ttgattttggagggatctcg-3′ |
| *gapdh* (mouse) | 5′-accacagtccatgccatcac-3′ | 5′-tccaccaccctgttgctgta-3′ |
| *NLuc* | 5'-gtccgtaactccgatccaaag-3′ | 5′-tgccatagtgcaggatcacct-3′ |
| FLuc | 5′-gtgacttcccatttgccacc-3′ | 5′-tgatctggttgccgaagatg-3′ |
| FLuc (spliced mRNA) | 5′-gaggtgcgtcaaacagcgac-3′ | 5′-tttggcatcttccctcgagg-3′ |

Table S2. RNA probes used in RNA EMSA.

| probe | RNA oligonucleotide sequence |
|---|---|
| (G4C2)3 | 5′-GGGGCCGGGGCCGGGGCC |
| (G4C2)4 | 5′-GGGGCCGGGGCCGGGGCCGGGGCC-3′ |
| (G2C4)4 | 5′-GGCCCCGGCCCCGGCCCCGGCCCC-3′ |
| (CUG)8 | 5′-CUGCUGCUGCUGCUGCUGCUGCUG-3′ |
| MUT | 5′-GAGGCCGGGACCGAGACCGAGGCC-3′ |
| (A4C2)3 | 5′-AAAACCAAAACCAAAACC-3′ |
| ORN | 5′-AGGGUUAGGGUUAGGGUUAGGG-3′ |
| TERRA | 5′-UUAGGGUUAGGGUUAGGGUUAGGG-3′ |
| mTERRA | 5′-UUAGGGUUAGUGUUAGUGUUAGGG-3′ |
| GC stem-loop | 5′-GCGCGCGCGCGCGCGCGCGAGAGCGCGCGCGCGCGCGC-3′ |
| NEAT1 | 5′-GGGAGGGAGGGAGGG-3′ |
